# Alternative oxidase expression in mtDNA mutator mice improves blood phenotype but enhances inflammatory and stress responses in skeletal muscle

**DOI:** 10.1101/2023.03.03.530968

**Authors:** Lilli Pihlajamäki, Sini Pirnes-Karhu, Howard T. Jacobs, Marten Szibor, Anu Suomalainen

**Affiliations:** Stem Cells and Metabolism Research Program (STEMM), Faculty of Medicine, University of Helsinki, 00290 Helsinki, Finland; Research Program for Clinical and Molecular Metabolism (CAMM), Faculty of Medicine, University of Helsinki, 00290 Helsinki, Finland; Faculty of Medicine and Health Technology, 33014 Tampere University, Finland; Department of Cardiothoracic Surgery, Center for Sepsis Control and Care (CSCC), Jena University Hospital, Friedrich-Schiller University of Jena, Am Klinikum 1, 07747 Jena, Germany; Helsinki University Hospital, HUSLAB, 00290 Helsinki, Finland

## Abstract

Energetic insufficiency, excess production of reactive oxygen species (ROS) and aberrant signalling partially account for the diverse pathology of mitochondrial diseases. Whether interventions affecting ROS, a regulator of stem-cell pools, could modify somatic stem-cell homeostasis remains unknown. Previous data from mitochondrial DNA (mtDNA) mutator mice showed that increased ROS leads to oxidative damage in erythroid progenitors, causing lifespan-limiting anemia. Also unclear is how ROS-targeted interventions affect terminally differentiated tissues. Here, we set out to test in mtDNA mutator mice how ubiquitous expression of the *Ciona intestinalis* alternative oxidase (AOX), which attenuates ROS production, affects murine stem-cell pools. We found that AOX does not affect neural stem cells but delays the progression of mutator-driven anemia. Furthermore, when combined with the mutator, AOX potentiates mitochondrial stress and inflammatory responses in skeletal muscle. These differential cell-type-specific findings demonstrate that AOX expression is not a global panacea for the cure of mitochondrial dysfunction. ROS attenuation needs to be carefully studied regarding specific underlying defects before AOX can be safely used in therapy.

**Graphical abstract:** 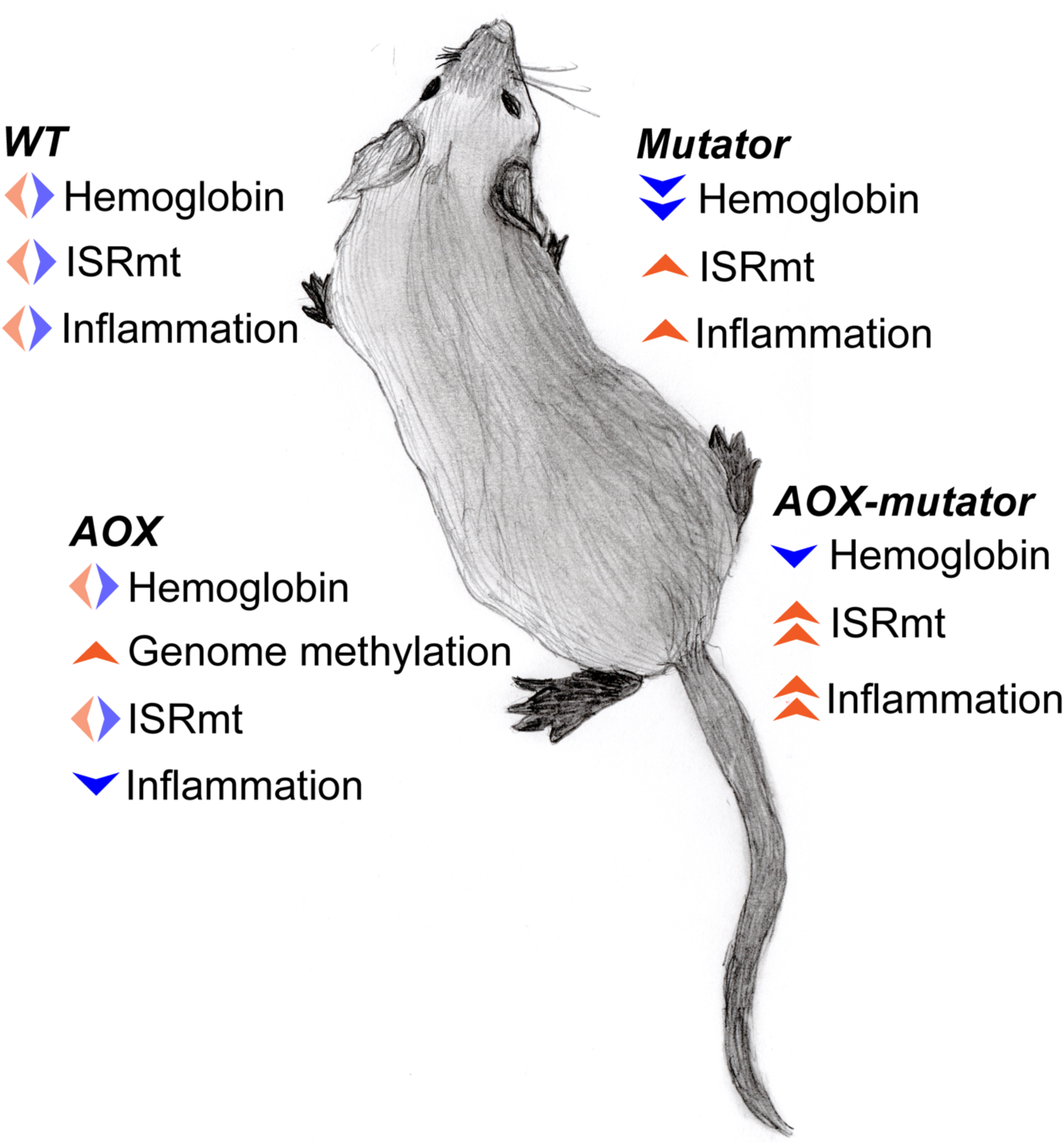

## Introduction

Genetic mutations that disrupt mitochondrial gene expression can lead to the defective assembly of respiratory chain (RC) complexes and appear to be a frequent cause of mitochondrial diseases. Myopathies, muscle diseases, are the most common mitochondrial disease phenotype in adults (1–3). Exactly how mitochondrial dysfunction, particularly RC dysfunction, leads to highly variable mitochondrial disease phenotypes is still poorly understood. Expanding this knowledge further is necessary to develop treatment options for these currently incurable diseases.

“Mutator” mice are a commonly used model to study mitochondrial dysfunction (4, 5). They carry a knock-in inactivating mutation in the exonuclease domain of the catalytic subunit of mitochondrial DNA polymerase gamma (POLG; p. D257A), leading to mtDNA mutation accumulation and increased mtDNA replication (6, 7).

These irregularities cause various cell-type-specific defects, especially affecting stem cell pools via increased ROS-related signaling (8–10). Furthermore, they show abnormal cell cycle progression and nuclear genomic DNA breakage in stem cells due to imbalanced nucleotide pools (7). These defects lead to a progeroid phenotype starting from 6–8 months of age with progressive hair graying, hair loss, osteoporosis, general wasting, and decreased fertility (4, 5). The lifespan of mutator mice is limited to 13–15 months by severe anemia, which develops alongside a decline in lymphopoiesis (11). In addition, postmitotic tissues demonstrate mildly progressive RC dysfunction, causing mild mitochondrial myopathy in skeletal muscle and cardiomyopathy (4, 5, 12, 13).

The excess production of reactive oxygen species (ROS) partially causes the mutator stem-cell homeostatic defect. In adult mutators, bone marrow shows highly increased oxidative stress within mitochondria, delayed mitochondrial exclusion from red blood cell (RBC) precursors, and extended iron loading by transferrin, leading to excess ROS generation via the Fenton reaction and oxidative damage in erythrocyte membranes (9). In other tissues, signs of ROS-related damage have not been found (4, 5). The antioxidant and reducing agent N-acetyl-L-cysteine (NAC) improves aberrant ROS signaling rescuing fetal neural stem-cell stemness and hematopoietic progenitor differentiation. In contrast, mitochondrially targeted ubiquinone, MitoQ, improves erythroid differentiation but is highly toxic to neural stem cells. (8, 10) These findings indicate differential sensitivities of stem-cell compartments to ROS-modifying treatments and highlight the importance of *in vivo* studies to collect evidence of cell-type-specific effects of mitochondrial-targeted interventions.

Alternative oxidase (AOX) resides in the inner mitochondrial membrane as part of the RC (Figure 1A) and is present in most eukaryotes, including metazoan taxa, but absent from insects and mammals. Under specific stress conditions, AOX expression mitigates over-reduction of the quinone pool and the consequent production of excess ROS, and the accumulation of keto acids (14, 15). In addition, AOX prevents the conversion of excess reducing power accumulated in the ubiquinone pool to superoxide. It does this by reducing O_2_ to H_2_O, bypassing the enzymes ubiquinol:cytochrome c oxidoreductase and cytochrome c oxidase (COX), i.e., complexes III and IV of the respiratory chain, respectively, and decreasing mitochondrial membrane potential and ATP synthesis (Figure 1A). This relaxes the coupling of electron transfer to ATP synthesis and decreases ROS production and reductive stress (Figure 1A). AOX from *Ciona intestinalis*, a sea squirt, has been xenotopically expressed in mammalian cells, flies, and mice (14–17). As proof of principle, AOX rescued fruit flies from death upon exposure to cyanide, an inhibitor of complex IV. AOX expression has been reported to be innocuous in human cell culture, *Drosophila*, and mice, except for one study, where AOX expression in a COX15 knockout mouse model worsened the myopathy and shortened the lifespan of these animals (14, 16, 18–24). AOX also compensates respiratory chain dysfunction, particularly when complex III or IV activity is limiting (14, 16, 18, 20). Different mouse tissues, including skeletal and heart muscle and blood cells, widely express AOX when driven by the synthetic CAG promoter (17). However, it is not known whether AOX is expressed in somatic stem cells and whether its effect on redox reactions affects stem-cell homeostasis.

**Figure 1.**
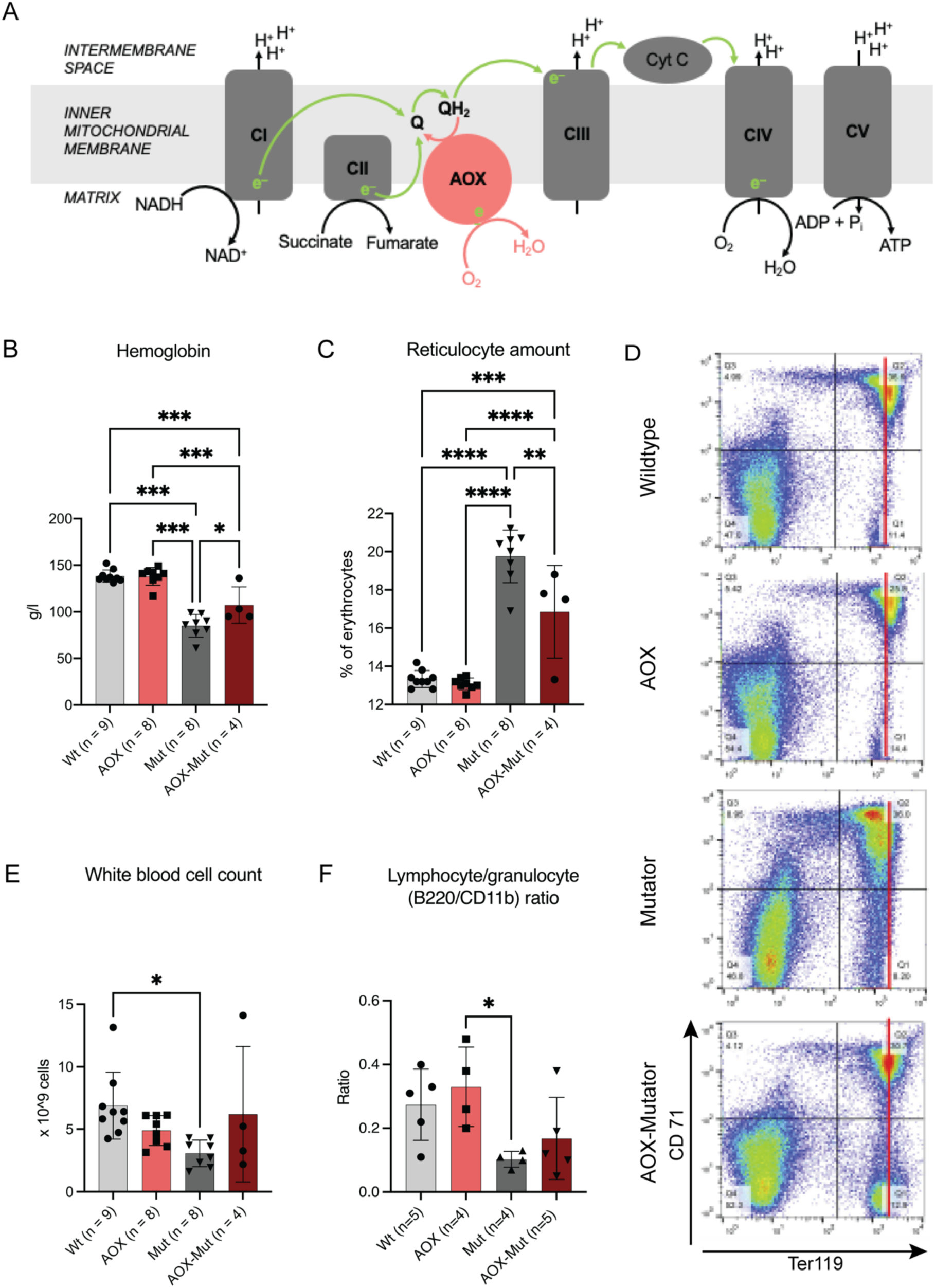
AOX expression alleviates anemia in 10-month-old mutators by shifting erythroid precursors toward a more mature state. A) A simplified scheme of the respiratory chain demonstrating the capacity of the alternative oxidase (AOX) to bypass functions of complex III (CIII) and IV (CIV). B–C) Hemoglobin and reticulocyte amounts from mouse blood. D) Hematopoietic precursor analysis from bone marrow. Erythrocyte FACS dot plots for Ter119 (maturing forms) and CD71 (early precursors). E) White blood cell count from mouse blood. F) Lymphocyte / granulocyte ratio calculated from fluorescence-activated cell sorting (FACS) data. Samples are biological replicates in the numbers presented in the figure; each sample was analyzed once. All graphs are mean with standard deviation (SD). Statistical significance determined using one-way ANOVA with p-values: * (p ≤ 0.05), ** (p ≤ 0.01), *** (p ≤ 0.001) and **** (p ≤ 0.0001). Abbreviations: CI–V, respiratory chain complexes; NADH, NAD^+^, reduced and oxidized forms of nicotinamide adenine dinucleotide; QH_2_, Q, reduced and oxidized forms of ubiquinone; WT, wildtype mice; AOX, AOX mice; AOX-Mut, AOX-Mutator mice; Mut, mutator mice.

In this study, we used the combination of AOX and mutator mice to determine whether and eventually how AOX expression affects a) somatic stem-cell function in healthy mice, b) stem cells subjected to mtDNA mutagenesis, and c) respiratory chain function in key post-mitotic tissues including skeletal muscle. We report that AOX expression has tissue-specific consequences in mutator mice. It alleviates anemia in mutators by shifting erythroid precursors toward a more mature state in bone marrow while also inducing the mitochondrial integrated stress response (ISRmt) and inflammatory pathways in skeletal muscle. Our findings indicate that AOX expression may be helpful in hematopoietic stem cells experiencing mitochondrial stress, but potentially harmful in postmitotic tissues via the induction or exacerbation of metabolic stress and inflammation.

## Results and Discussion

### Generation of AOX-mutator mice

The mouse strains used in the study, i.e., those expressing *Ciona intestinalis* AOX (AOX mice) and those with a knock-in mutation (p.D257A) inactivating the proof-reading exonuclease domain of DNA polymerase gamma (POLG; mutator mice) have been described previously in (17) and (4, 5), respectively. The strains were maintained in the C57Bl6/JOlaHsd background, AOX mice as hemizygotes and mutators as heterozygotes. The mutator allele was transferred only via the paternal line, to prevent the accumulation of maternally inherited mtDNA mutations. We obtained double-transgenic mice expressing AOX that were also homozygous for the POLG (p. D257A) mutator mutation. AOX, mutator, AOX-mutator, and wildtype (WT) littermates were born approximately in the expected Mendelian ratios.

The gross phenotype and outward appearance of AOX-mutator double-transgenic mice was indistinguishable from that of mutator mice, with signs of progeria starting from the age of 6 months. Neural stem cells (NSCs) and postmitotic skeletal and heart muscle were confirmed to express AOX (Figure S1A–C).

### AOX expression alleviates defective hematopoiesis in mutators

Mutator mice manifest anemia and increased amounts of reticulocytes (4, 8, 11). Therefore, we asked whether the effect of AOX on redox metabolism affects hematopoietic progenitor differentiation in WT or mtDNA mutator mice. We harvested bone marrow and peripheral blood of WT and mutator mice with and without AOX expression at 43 weeks. At this age, mutators show various progeroid signs: kyphosis, alopecia, weight loss, osteoporosis, and anemia.

AOX-mutators showed higher hemoglobin (Hb) and lower circulating reticulocyte (immature RBC) counts than mutators (Figure 1B–C). AOX expression did not affect the number of erythroid precursors in adult mutator bone marrow (Figure S2A–B). However, fluorescence-activated cell sorting (FACS) analysis revealed that mutators show lower mean intensity of the Ter119 signal and an absence of cells with selective high Ter119, whilst AOX-mutator bone marrow shows Ter119 signal with mean intensity resembling WT controls and the presence of cells with Ter119 high only, as seen in WT (Figure 1D, Figure S2A–B). These results show that AOX expression shifts the maturation pattern of mutator erythroid precursors towards that seen in WT. Total white blood cell counts trended towards WT values in AOX-mutators compared to mutators (Figure 1E), as did bone marrow lymphocyte/granulocyte ratios (Figure 1F). However, the variation in these counts was considerable in all groups (Figure 1E–F). We observe that AOX partially rescues the abnormal erythrocyte differentiation pattern seen in mutators (Figure 1D, Figure S2A–B) similarly to NAC (8) and MitoQ (10), a modified ubiquinone targeted to accumulate in mitochondria (25). In conclusion, AOX affects erythrocyte differentiation in mutators but not in WT mice and partially rescues the anemia, which is ultimately fatal to mutator mice.

Because of the effect of AOX on mutator hematopoiesis, we proceeded to look at its effect on neural stem cells (NSCs). Previously, we demonstrated that mutator NSCs show decreased “stemness”, presenting a decreased ability to self-renew in clonal culture (8). AOX expression did not rescue defective self-renewal or proliferation of mutator NSCs (Figure S2C–D). Compared with erythrocyte precursors, mutator NSCs appear to be less sensitive to the effects of antioxidants, with only NAC treatment, but neither MitoQ nor AOX able to restore their self-renewal capacity (14). AOX expression did not affect the stemness or growth of WT NSCs (Figure S2C–D). While AOX is expressed in most tissues, including NSCs (Figure S1A) (17), differential expression of AOX in bone marrow or in different blood-cell populations has not been studied, and might explain the distinct effects on neural progenitors and hematopoietic cells.

These data demonstrate differential effects of AOX expression on different somatic stem-cell compartments, but only under conditions of mitochondrial dysfunction. In WT mice AOX expression had no effect on erythroid, lymphatic, or neural progenitors or their differentiation pattern.

### AOX expression affects respiratory chain components in mutator skeletal muscle

After examining the effect of AOX on stem- and progenitor-cell compartments, we proceeded to ask whether it could provide a therapeutic benefit and alleviate RC dysfunction previously demonstrated in the post-mitotic skeletal muscle of mutators (4, 8). Histochemistry for complex IV (COX, partially encoded by mtDNA) and succinate dehydrogenase (SDH, complex II, nuclear-encoded) revealed no COX-negative, SDH-positive fibers indicative of mitochondrial dysfunction in the skeletal muscle of mice of any of the genotypes analyzed (Figure 2A). However, overall COX staining intensity was decreased, compared with WT, both in mutators and AOX-mutators, consistent with diminished COX activity (Figure 2C). Only in mutators we observed a subset of COX muscle fibers with strong staining, from here referred to as “hyperpositive”, a novel finding (Figure 2A–B). AOX-mutator skeletal muscle was lacking in these highly COX-positive fibers but had a similar overall COX staining intensity (Figure 2A–C). Increased COX activity is indicative of increased mitochondrial activity; AOX is known to decrease mitochondrial biogenesis upon mitochondrial dysfunction in COX15-deficient mice (22). Therefore, the expression of AOX does not rescue COX protein amounts to the wildtype level (Figure 2C, Figure S3G), but seems to impact the dysregulation of RC function in the muscle of mutators.

**Figure 2.**
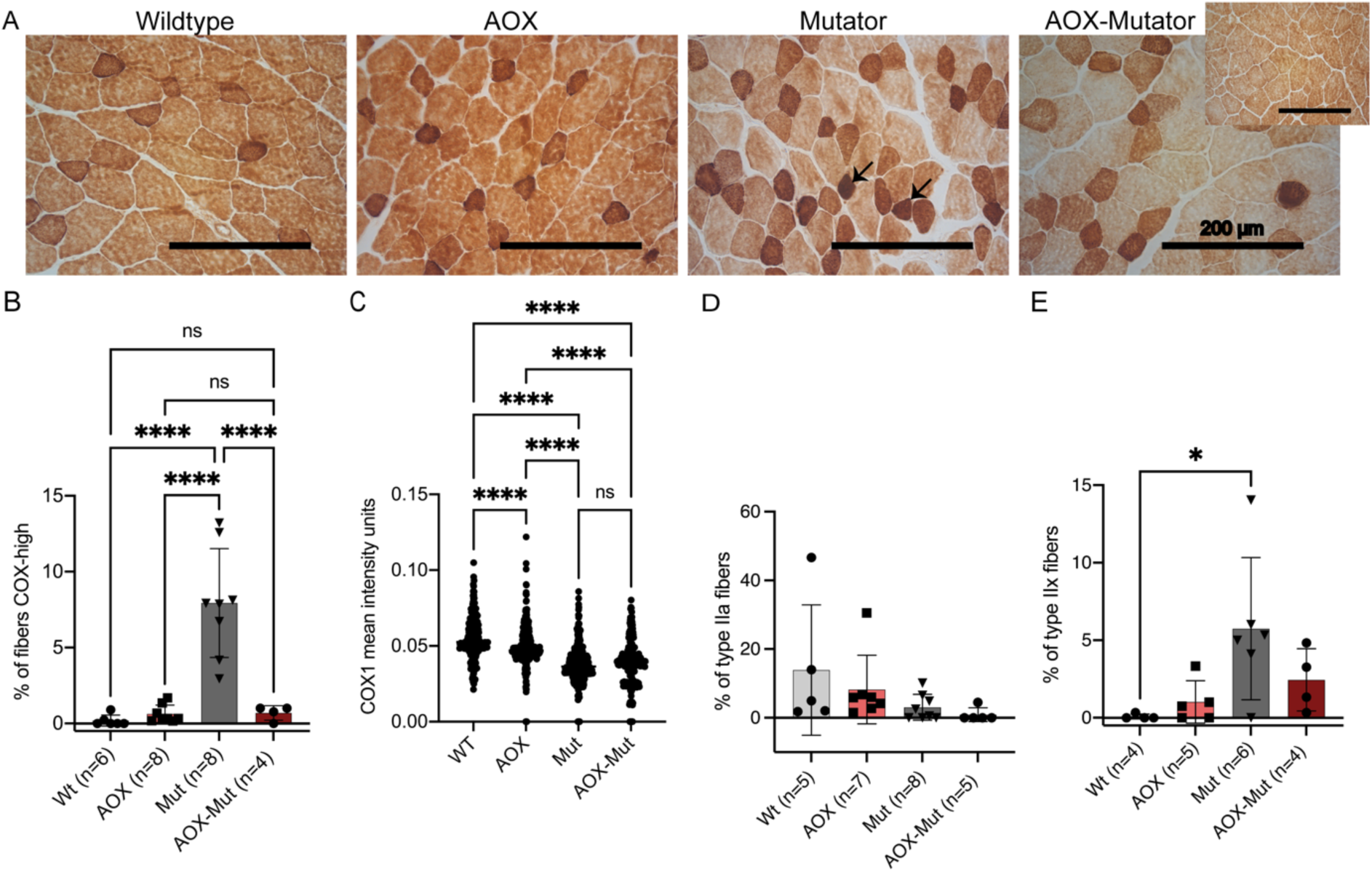
AOX expression affects respiratory chain components in Mutator skeletal muscle. A) Cytochrome *c* oxidase (COX, brown precipitate) and succinate dehydrogenase (SDH, blue precipitate; visible only in cells with COX deficiency) histochemical activity assay from skeletal muscle of wildtype, AOX, mutator and AOX-Mutator mice. Small image for AOX-Mutator shows homogeneous staining seen commonly in some sections. Arrows indicate example fibers with intense COX activity. Magnification 20x, scale bar 200 µm. B) Quantification of fibers with intense COX activity; COX-SDH histochemical activity assay on frozen sections of skeletal muscle. (C) Scatterplot of COX1 mean intensity units; IF COX1 analysis of (B) in skeletal muscle. (D–E) Quantification of muscle fiber types, IF analysis of myosin heavy chain (MHC) type IIa (D) and type IIx (E) in skeletal muscle. Samples are biological replicates in the numbers presented in the figure; each sample was analyzed once. All graphs are mean with standard deviation (SD). Statistical significance determined using one-way ANOVA with p-values: * (p ≤ 0.05), ** (p ≤ 0.01), *** (p ≤ 0.001) and **** (p ≤ 0.0001).

We proceeded to look at RC components at the protein level, by immunofluorescent staining for the COX1 subunit, which was also decreased in mutators and not rescued by AOX (Figure 2C, Figure S3A). AOX also did not affect COX1 expression in otherwise wildtype mice (Figure 2C, Figure S3A). Immunohistochemistry for the SDHA subunit of the fully nuclear-coded complex II showed no difference between the groups (Fig. S3B–C).

We also asked whether the alteration in COX activity and protein amount in mutators could be related to a shift in muscle fiber type. We explored myosin heavy chain (MHC) isoforms preferentially associated with either oxidative or glycolytic metabolism in skeletal muscle. Based on immunohistochemistry for myosin heavy chain (MHC) expression, which enables different muscle fiber types to be distinguished (26, 27), mutators showed a decreased representation of smaller, oxidative type IIa fibers compared to wildtype mice (Figure 2D, Figure S3D), but this was unaffected by AOX expression (Figure 2D, Figure S3D). Glycolytic muscle fibers were prevalent in all mouse groups (Figure S2E). However, mutators showed an increase in type IIx glycolytic fibers compared to the other groups (Figure 2E, Figure S2F). The shift toward glycolytic muscle metabolism is consistent with decreased oxidative phosphorylation (OXPHOS) and COX activity, although the mitigation of this effect in the presence of AOX suggests that AOX may interfere with the stress signaling involved.

In deletor mice, a well-characterized model of mitochondrial myopathy, RC deficiency manifests in a mosaic pattern in skeletal muscle, alongside mosaicism for the promotion or prevention of mitophagy (28). Our finding of COX-hyperpositive fibers in mutator skeletal muscle further highlights the need for single-cell studies in the pathology of mitochondrial dysfunction, since a finding of decreased average COX levels would overlook both types of mosaicism.

In the brain, aging mutator mice show diminished COX activity in multiple brain regions (cortex, hippocampus, accumbens, striatum, and thalamus) with a smaller decrease in the cerebral cortex (29, 30). In agreement with our previous results (8), we were unable to identify COX-negative, SDH-positive neurons in the hippocampus or dentate nucleus of mutators. However, RC-deficient cells were present around the lateral ventricles in the subventricular zone where NSCs reside. AOX expression did not alter the pattern of COX-SDH activity staining in the brain (Figure S3H–I).

### AOX expression in wildtype mice leads to altered genome methylation and decreased inflammation

In plants, AOX modifies mitochondrial signaling and affects nuclear transcription programs (31). To gain insight into the effects of AOX in mice, we performed RNA sequencing (RNAseq) on mouse skeletal muscle with or without AOX, in the WT and mutator backgrounds (Figure 3). Two principal components explained 58% of the variance between samples (Figure 3A). The number of transcripts significantly differing between mutator or AOX-mutator and wildtype (Figure 3F, 3H) was substantially higher than between AOX and wild-type (Figure 3D) or AOX-mutator and mutator (Figure 4A, Dataset S1).

**Figure 3.**
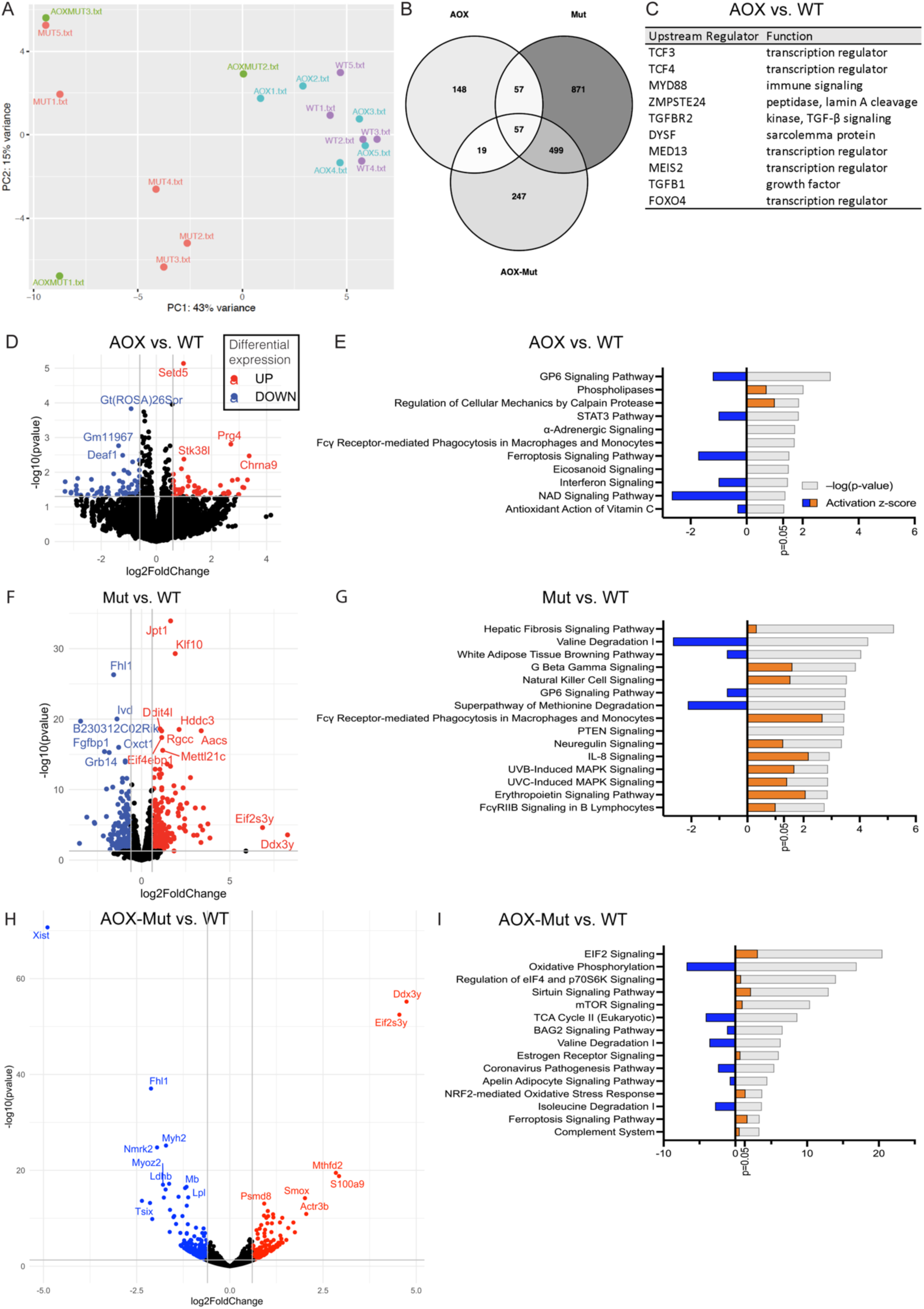
AOX and mutator mice have distinct gene expression patterns in skeletal muscle. A) Factors explaining variance in RNA sequencing (RNA-seq) of skeletal muscle samples analyzed using principal components analysis. B) Overlap of significantly changed transcripts in RNA-seq in AOX, Mut and AOX-Mut compared to WT. C) Top 10 activated upstream regulators in AOX compared to WT in RNA-seq data of mouse skeletal muscle analyzed using Ingenuity Pathway Analysis (IPA). D, F, H) Transcripts with highest experimental fold change differing significantly between D) AOX, F) Mut, and H) AOX-Mut vs. WT in RNA-seq data of mouse skeletal muscle shown as volcano plots. E, G, I) Comparison of the most significant canonical pathways changed in E) AOX, G) Mut, I) AOX-Mut vs. WT based on RNA-seq data of mouse skeletal muscle analyzed using IPA. WT (n=5), AOX (n=5), Mutator (n=5), and AOX-Mutator (n=3), biological replicates, RNA-seq for each sample was performed once. Abbreviations: WT, wildtype mice; AOX, AOX mice; AOX-Mut, AOX-Mutator mice; Mut, mutator mice.

**Figure 4.**
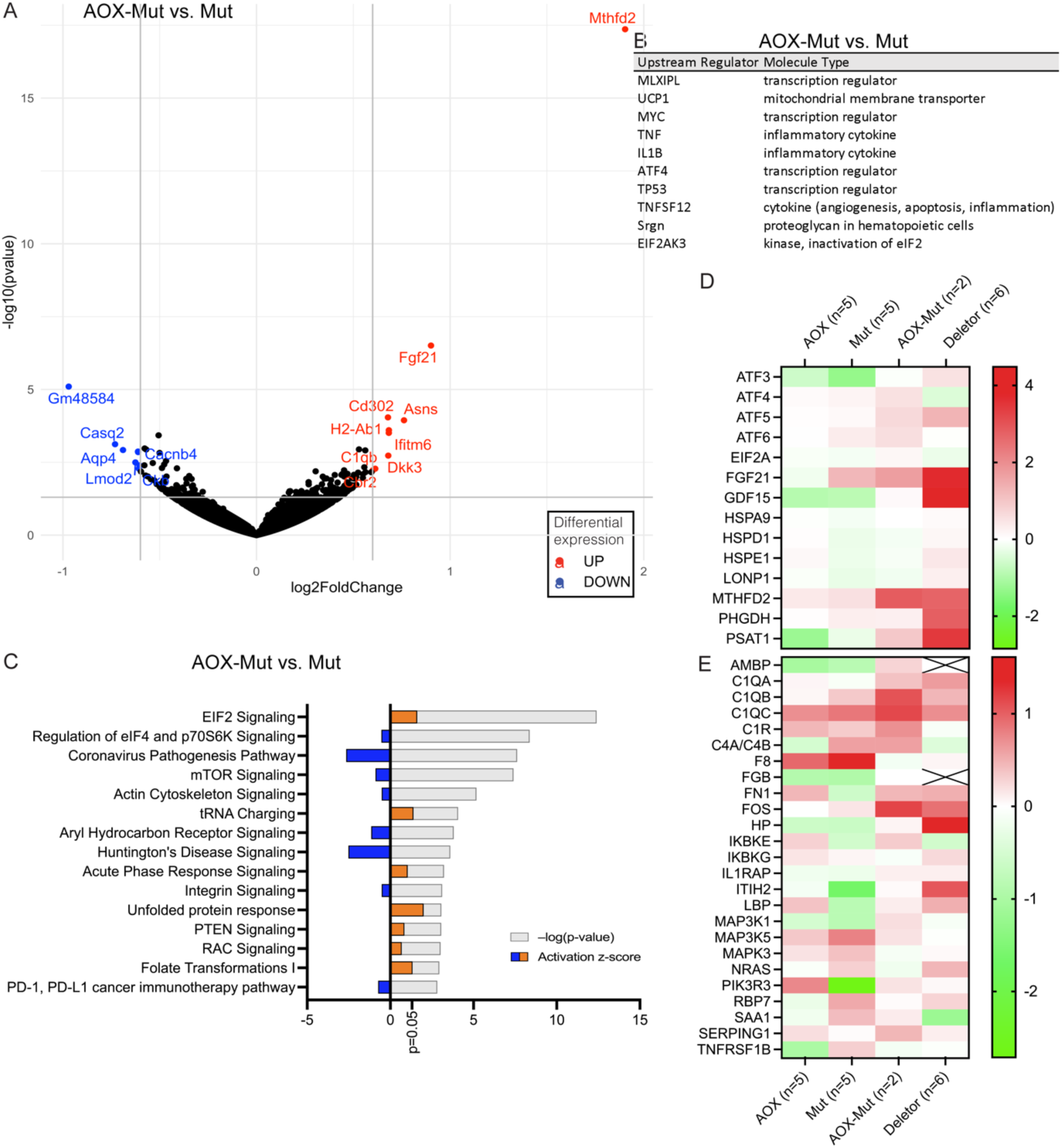
AOX expression on a mutator background leads to activation of the mitochondrial integrated stress response (ISRmt) and inflammatory pathways. A) Transcripts with highest experimental fold change differing significantly in AOX-Mut vs. Mut in RNA-seq data of mouse skeletal muscle shown as a volcano plot. B) Top 10 activated upstream regulators in AOX-Mut compared to Mut in RNA-seq data of mouse skeletal muscle analyzed using IPA. C) Comparison of the most significant canonical pathways changed in AOX-Mut vs. Mut based on RNA-seq data of mouse skeletal muscle analyzed using IPA. D–E) Heatmaps of RNA-seq data of mouse skeletal muscle analyzed using IPA showing D) ISRmt genes and E) acute phase response genes. WT (n=5), AOX (n=5), Mutator (n=5), and AOX-Mutator (n=3), biological replicates, RNA-seq for each sample was performed once. Abbreviations: WT, wildtype mice; AOX, AOX mice; AOX-Mut, AOX-Mutator mice; Mut, mutator mice.

The histone methyltransferase Setd5 was the most significantly altered transcript in AOX vs. WT (Figure 3D), whilst many methylation-regulated genes (e.g., Chrna9, Prg4, Zbp1, and Zdbf2) were among those with the highest fold change (Figure 3D, Table S1). Posttranslational modifications such as methylation play an important role as determinants of innate immunity and inflammatory responses (33). Moreover, mitochondrial proteins, mtDNA, or RNA released into the cytoplasm, e.g., because of mitochondrial dysfunction, could act as a pro-inflammatory signal. The most activated upstream regulators in AOX vs. WT skeletal muscle are immune-related, notably transcription factors TCF 3 and 4 playing a role in lymphopoiesis, and myeloid differentiation primary response factor 88 (MyD88), involved in NFκB activation (Figure 3C). Overall, inflammatory and immune-related pathways, such as STAT3 and interferon signaling and phagocytosis, were downregulated in the skeletal muscle of AOX mice (Figure 3E–G). Other highly activated upstream regulators include dysferlin (DYSF), which is thought to be involved in muscle fiber repair (Figure 3C).

### AOX expression in mutator mice leads to activation of inflammatory pathways and ISRmt

In contrast to AOX mice, multiple immune-related pathways, including natural killer (NK) cell and IL-8 signaling and phagocytosis, were predicted to be upregulated in mutator skeletal muscle (Figure 3G). Furthermore, when AOX was also present, pro-inflammatory signaling was even more highly stimulated (Figure 4C), notably IL-1β-driven acute phase response signaling (32) (Figure 4B, C, E). Top upstream regulators in AOX-mutators compared to mutators included inflammatory cytokines (IL1β, TNF), and TP53, a major tumor suppressor (Figure 4B, Table S2).

When comparing AOX-mutator to the other mouse groups, molecules involved in signaling of the mitochondrial integrated stress response (ISRmt), namely MTHFD2, FGF21, PSAT1, PHGDH, and GDF15, a metabokine that mediates systemic ISRmt signaling alongside FGF21, were upregulated (Figure 4A, D, Table S1). At the regulator level, in AOX-mutators compared to mutators, IPA showed marked activation of ATF-regulated genes and the ISRmt-inducing (33) uncoupling protein 1 (UCP1), which dissipates heat by uncoupling the mitochondrial proton gradient from respiration (Figure 4B, Table S2). In contrast, ISRmt activation seemed to be independent of mTOR signaling, a known regulator (34) (Figure 4C), and of the newly described OMA1–DELE1–HRI regulatory pathway (35) (Dataset S1). Further, the unfolded protein response (UPR) and folate signaling – both components of ISRmt – were activated (Figure 4C). Upon AOX expression, a similar increase in FGF21 and GDF15 signaling, one-carbon signaling, and UPRmt independent of mTOR signaling was seen previously in a mouse model of mitochondrial myopathy highlighting potential risks of interfering with ROS signaling in mitochondrial myopathies (22).

While AOX expression has previously been thought to be mostly innocuous (14, 16, 18–21, 23), we show here that it induces stress and inflammatory responses in a postmitotic tissue experiencing a broad and progressive mitochondrial RC dysfunction, involving activation of the previously characterized ISRmt (34). Since AOX appears to attenuate rather than enhance ROS signaling (21, 22), our findings indicate that ISRmt induction can take place independently of a ROS signal.

The co-induction of ISRmt and of inflammatory signaling in AOX-mutator muscle may be independent or may be linked. Phosphorylation of eIF2α in response to a variety of physiological stresses, including infection, is an initiator of both processes (36–38). However, AOX limits an IL-1β-dependent response in bone-marrow-derived macrophages (24), suggesting that its effect(s) on inflammation may be tissue-specific. Further research is needed into the significance of ISRmt induction in a setting of mitochondrial dysfunction.

### AOX expression or the Polg mutator alters innate immune signaling

Given the upregulation of multiple pro-inflammatory mediators in AOX-mutator mice, we examined innate-immunity-related genes in our muscle RNA-seq data. Multiple regulators of innate immunity were upregulated in mutators vs. WT mice (Table S2), including IRF3, a critical player in the cGAS-STING pathway (39) which is triggered upon cytosolic sensing of mtDNA (40, 41). This has been suggested to contributed to pathologies involving mitochondrial dysfunction (42–47), and led us to look at the cGAS-STING pathway in more detail using our RNA-seq data.

cGAS and STING, as well as the cGAS-STING-TBK1-IRF3 signaling cascade, were upregulated by expression of AOX and/or the mutator phenotype (Figure 5A–B), and this was true also for deletor mice, a model for mitochondrial myopathy carrying a dominant patient-equivalent mutation in the mitochondrial replicative helicase Twinkle (48).

**Figure 5.**
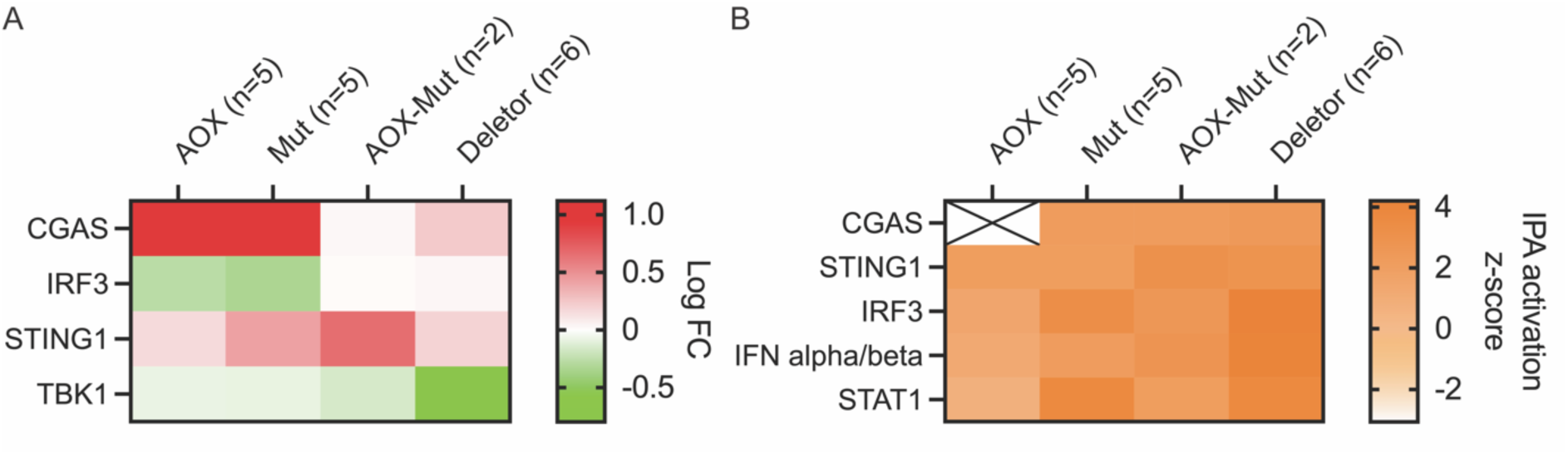
Activation of innate immunity pathways in Mutators is altered by AOX. Transcriptome analysis; effect of AOX on skeletal muscle. Heatmaps of cGAS-STING pathway genes; RNA-seq data, mouse skeletal muscle; analysis by Ingenuity Pathway Analysis showing comparison based on A) experimental logarithmic fold change (Log FC) and B) predicted activation state (z-score). Samples are biological replicates in the numbers presented in the figure; RNA-seq for each sample was performed once.

Aberrant activation of the type I IFN response may aggravate the mutator phenotype (49). This has also been shown in other cases of mitochondrial dysfunction, such as in TFAM (transcription factor A, mitochondrial) heterozygous knockout (*Tfam^+/-^*) mice (45) and in mice with mtDNA stress induced by exhaustive exercise or mtDNA mutations in the absence of parkin or PINK1 (44). Activation of innate immunity pathways may therefore be a common response to mitochondrial stress.

Despite much research, no specific therapy is available for pathological OXPHOS defects. AOX expression has been proposed as a tool not only to study mechanisms of respiratory chain dysfunction but also to alleviate its pathological consequences (15, 18). The present findings indicate that implementation of AOX, in combination with at least one type of mitochondrial impairment, namely the lifetime accumulation of mtDNA point-mutations, activates various stress and immune-related pathways in a tissue-specific fashion. These may be either beneficial, as is the case for the alleviation of mutator-driven anemia, or potentially harmful, as for inflammatory responses in skeletal muscle. Thus, to have any use in therapy, the obvious ethical problems associated with the use of genetic therapies in humans will need to be considered and the effects of AOX expression will need to be carefully studied in different tissues and in response to different mitochondrial defects. More broadly, the nature of the link between mitochondrial dysfunction, the stress responses that it induces, and altered innate immune signaling warrants wider investigation.

## Materials and Methods

### Mouse models

AOX mice were generated and characterized by Szibor et al. (17). Mutator mice (4, 5), also extensively characterized, were generously supplied by Tomas Prolla. Mutator mice are also commercially available (IMSR_JAX:017341, The Jackson Laboratory). Hemizygous AOX mice and heterozygous mutator mice were first crossed. The resultant AOX hemizygous – mutator heterozygous females were further crossed with a) heterozygous mutator males or b) homozygous mutator males. Using method a), the second breeding step resulted in AOX hemizygous – mutator heterozygous mice (5.6%, not used in the study) and heterozygous mutator mice (22.2%, not used in the study), as well as the study groups actually used: AOX (27.8%), mutator (22.2%), AOX-mutator double-transgenic (16.7%), and wildtype littermate mice (27.8%), close to expected Mendelian ratios (16.7% for each group). Using method b), the second breeding step resulted in AOX hemizygous – mutator heterozygous mice (20.0%, not used in the study) and heterozygous mutator mice (13.3%, not used in the study) as well as the study groups AOX-mutator double-transgenic (23.3%) and mutator mice (23.3%) in approximately Mendelian rations (25% for each group); age-matched WT and AOX pups were used. (Figure S4) Using methods a) and b), 46 mice were obtained. An additional 13 mice were used for NSC extraction. Mice of both sexes were used in the study. No animals were excluded from analyses. Randomization and blinding were not applicable in this study. The National Animal Experiment Board of Finland approved animal maintenance and experimentation (permit ESAVI-689-04.10.07-2015), and the mice were maintained and studied according to 3R principles. ARRIVE guidelines were followed as applicable.

### NSC

NSC extraction from WT (n=4), AOX (n=2), Mutator (n=4), and AOX-Mutator (n=3) mice’s embryos was performed from the lateral ventricular wall of E11.5–E15.5 mouse brain as previously described (50). Neurospheres were cultured in serum-free Ham’s F12 medium (Sigma-Aldrich, N4888) supplemented with B27 (Gibco, 12587010), GlutaMAX^TM^ (Gibco, 35050061), penicillin-streptomycin (Gibco, 15070073), FGF (Sigma-Aldrich, F0291), and EGF (BD, 354052). Cell cultures were tested monthly for mycoplasma contamination. Analysis of NSC self-renewal capacity was performed as previously described (50). To determine proliferation rate, a BrdU incorporation assay was used: neurospheres were incubated in 10 μM BrdU (BD PharMingen), stained with anti-BrdU and fluorescent secondary antibody, and analyzed using a FACSAria Cell Sorter.

### Western blot

Whole-cell protein extraction from NSCs was performed as in (8). Protein concentration was measured using the Bradford method (Protein Assay, Bio-Rad). For SDS-PAGE prior to Western blotting, sample aliquots were mixed with 3 × SDS loading buffer, denatured, and resolved on 4–20% Mini-PROTEAN® TGX Stain-Free^TM^ Gels (Bio-Rad, #4568096) at 100–120 V for 1 h. Proteins were transferred to PVDF membranes using Trans-Blot Turbo RTA Mini transfer kit (Bio-Rad, #1704727). Membranes were blocked with 5% milk in 1x TBST for 1 h at room temperature, then probed overnight at 4°C with a custom-manufactured anti-AOX antibody (polyclonal rabbit serum, 1:33000, (51)) in 5% milk, then washed three times in blocking solution. Incubation with HRP-conjugated goat anti-rabbit secondary antibody (Jackson ImmunoResearch, 111-035-144, 1:10000) was for 1 h at room temperature followed by three washes in TBST. Imaging was done using a ChemiDoc imaging system (Bio-Rad).

### Blood analyses

Blood was collected from sacrificed mice into EDTA tubes (Greiner Bio-One) by heart puncture. Blood count was performed from whole blood samples on the same day using an Advia 2120i analyzer (Siemens).

### FACS

The femoral and tibial bones of sacrificed mice were cut out and cleaned of soft tissue. Bone marrow was flushed out using a syringe and 1 ml of PBS + 5% FBS and filtered through a 40 μm strainer (Dako). Cells were counted and analyzed immediately. Hematopoietic lineages of adult bone marrows were analyzed using a BD Influx Cell Sorter (Beckton Dickinson). CD16/CD32 blocker (BD Pharmingen, 1 μg/1 × 10^6^ cells) was used to inhibit possible nonspecific binding of the antibodies. Fluorescence-conjugated antibodies against CD71 (BD Pharmingen), CD11b (BD Pharmingen), Ter119 (eBioscience) and B220 (BD Pharmingen) were used (each 1 μg / 1×10^6^ cells) for staining. Propidium iodide was used as a dead cell marker. Unstained and fluorescence-minus-one (FMO) controls were used to determine the autofluorescence of the cells and gates for the cell populations of interest, respectively. One hundred thousand cells per sample were analyzed.

### RNA extraction

Total RNA was extracted from skeletal muscle (*m. quadriceps femoris*) using TRIzol reagent (Invitrogen) and purified using RNeasy Mini Kit (Qiagen). The samples were homogenized using TRIzol reagent (Invitrogen), and chloroform was then added to allow separation of the homogenate. Lysate-ethanol mix was then purified using RNeasy Mini Kit (Qiagen) according to the manufacturer’s instructions. Extracted RNA was treated with RNase-free DNAse (M6101, Promega). 1000 ng of total RNA was used to generate cDNA using MAXIMA cDNA synthesis kit (K1641; Thermo Scientific).

### Transcriptomics analysis

RNA from WT (n=5), AOX (n=5), Mutator (n=5), and AOX-Mutator (n=3) was submitted for transcriptomic analysis. RNA quality control analysis was done using Tapestation 4200 (Agilent). RNA sequencing was performed using a “Bulkseq” 3’UTR-counting gene expression profiling method based broadly on BRBseq/Dropseq with high output (1×75 bp) read lengths. The service was provided by the Biomedicum Functional Genomics Unit at the Helsinki Institute of Life Science and Biocenter Finland at the University of Helsinki. Primary data analysis was done using the DeSeq2 package from Bioconductor release 3.9 (53) in R Studio version 1.4.1103 using R version 3.6.3. Data were analyzed further through IPA (QIAGEN Inc., https://www.qiagenbioinformatics.com/products/ingenuity-pathway-analysis) (54).

### Histology

Skeletal muscle (*m. quadriceps femoris, QF*), heart muscle, and brain were collected from wildtype, AOX, Mutator, and AOX-mutator mice aged 40 weeks. The tissues were harvested immediately after sacrificing the mice, embedded in OCT Compound Embedding Medium (Tissue-Tek), and snap-frozen in 2-methylbutane bath in liquid nitrogen. In situ histochemical COX and SDH activities were analyzed from frozen tissue sections (12 μm) using standard protocols (48). Imaging was done by light microscopy (Axioplan 2 Universal Microscope, Zeiss). Approximately 200 – 700 fibers from each mouse in the study groups WT (n=6), AOX (n=8), Mutator (n=8), and AOX-Mutator (n=4) were counted to calculate the percentage of COX-negative and COX-negative/SDH-positive fibers from QF sections. The COX-negative and SDH-positive fibers were quantified from each study group using ImageJ software (55).

Immunofluorescent staining of SDHA (complex II), COX-I (complex IV), nuclei, and laminin in muscle of WT (n=3), AOX (n=3), Mutator (n=4), and AOX-Mutator (n=2) mice was done by adapting a previously published protocol (56). Primary antibodies used were anti-SDHA mouse IgG1 (Abcam, ab14715, 1:95), anti-MTCO1 mouse IgG2a (Abcam, ab14705, 1:95), and anti-laminin (Sigma-Aldrich L9393, 1:100). Secondary antibodies used were goat anti-mouse IgG1 biotin (Abcam, ab97238, 1:200) with streptavidin conjugate Alexa Fluor 647 (Invitrogen, S21374, 1:100), goat anti-mouse IgG2a Alexa Fluor 488 (Invitrogen, A-21136, 1:200), and goat anti-rabbit Alexa Fluor 568 (Invitrogen, A-11011, 1:100), respectively. Slides were mounted with ProLong Diamond Antifade Mountant with DAPI (Invitrogen, P36966). Images were generated using 3DHISTECH Pannoramic 250 FLASH II digital slide scanner at Genome Biology Unit supported by HiLIFE and the Faculty of Medicine, University of Helsinki, and Biocenter Finland. Digitalized images were visualized using 3DHISTECH CaseViewer 2.2.0 software. Quantification of SDHA and COX-I staining intensity and fiber size was done using CellProfiler (57, 58).

Myosin heavy chain (MyHC) isoforms were stained in skeletal muscle (*m. quadriceps femoris*) of WT (n=5), AOX (n=7), Mutator (n=8), and AOX-Mutator (n=5) mice using a protocol adapted from previous reports (59, 60). Fresh frozen 10 μm muscle sections were used. Primary antibodies were obtained from the Developmental Studies Hybridoma Bank at the University of Iowa: MyHC I antibody BA-D5 1:40 (61), MyHC IIa antibody SC-71 1:200 (61), MyHC IIb antibody BF-F3 1:100 (62), and MyHC IIx antibody 6H1 1:50 (63). Secondary antibodies used were goat anti-mouse IgG2a Alexa Fluor 488 (Invitrogen, A-21131, 1:200) with BA-D5, goat anti-mouse IgG (H+L) Alexa Fluor 488 (Invitrogen, A-11001, 1:200) with SC-71, and goat anti-mouse IgM Alexa Fluor 488 (Invitrogen, A-21042, 1:100) with BF-F3 and 6H1. Additionally, slides were co-stained for laminin (primary antibody: Sigma-Aldrich, L9393, batch 82508, 1:100) with secondary antibody goat anti-rabbit IgG (H+L) Alexa Fluor 594 (Invitrogen, R37119, 1:300). Slides were mounted with Vectashield® Antifade Mounting Medium with DAPI (H-1200-10, Vector). Images were taken using the Axio Imager M1 microscope (ZEISSAXIOM1, ZEISS). Quantification of positive fibers for each MyHC type was done using ImageJ software (55).

### Statistical analyses

One-way ANOVA for groups displaying normal distribution followed by Tukey’s multiple comparisons test was performed on independent observations using GraphPad Prism version 9.0.2 for Mac, GraphPad Software, San Diego, USA, www.graphpad.com. P-values are shown as ns (p > 0.05), * (p ≤ 0.05), ** (p ≤ 0.01), *** (p ≤ 0.001), and **** (p ≤ 0.0001). Graphs show the mean with standard deviation (SD) with individual measurements shown.

## Acknowledgements

We are grateful to Markus Innilä, Babette Hollman, Sonja Jansson, Maarit Partanen, and Tea Tuomela for technical assistance; Mito Takayuki, Christopher Jackson, and Nahid Khan for assistance with imaging; Gulayse Ince-Dunn for RNAseq insight; Yilin Kang for helpful discussions; and Biomedicum Imaging Unit and Biomedicum Functional Genomics Unit for services and infrastructure. The authors acknowledge funding from Biomedicum Helsinki foundation (LP), The Finnish Medical Foundation (LP), Academy of Finland (AS, HTJ, MSz), Sigrid Jusélius foundation (LP, AS), the University of Helsinki (AS), and the European Research Council (HTJ, MSz). The work used core facilities part-supported by funding from Biocenter Finland.

---

**Figure S1.**
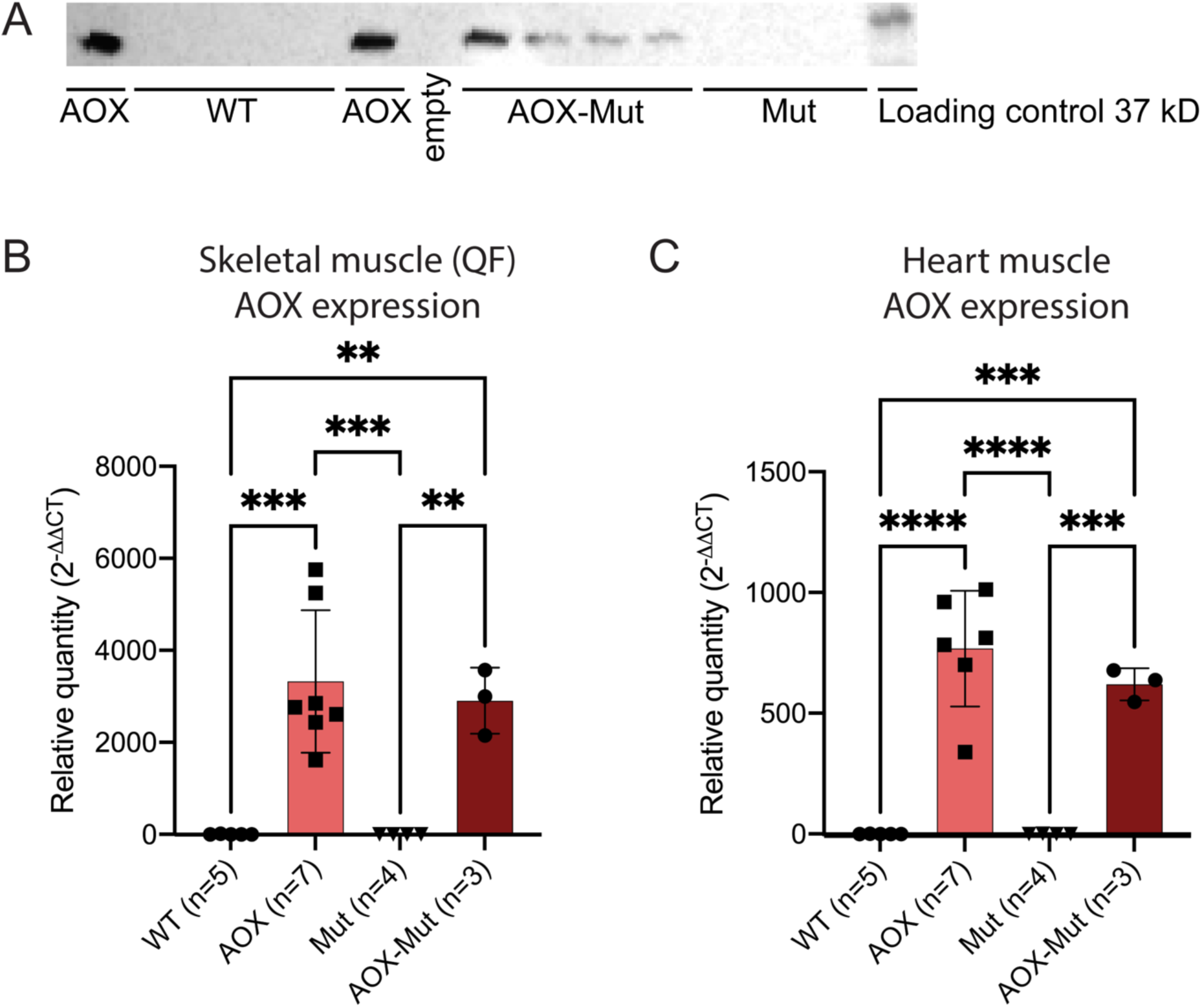
Expression of AOX in different tissues. A) AOX expression in neural stem cells (NSCs), western blot. B–C) AOX expression in skeletal (B) and heart (C) muscle of WT, AOX, Mut, and AOX-Mut mice as measured by quantitative RT-PCR. Samples are biological replicates in the numbers presented in the figure; each sample was analyzed once. All graphs are mean with standard deviation (SD) with values for individual mice shown. Statistical significance determined using one-way ANOVA with p-values: * (p ≤ 0.05), ** (p ≤ 0.01), *** (p ≤ 0.001) and **** (p ≤ 0.0001). Samples are biological replicates in the numbers presented in the figure; each sample was analyzed as three technical replicates. Abbreviations: WT, wildtype mice; AOX, AOX mice; Mut, mutator mice; AOX-Mut, AOX-Mutator mice.

**Figure S2.**
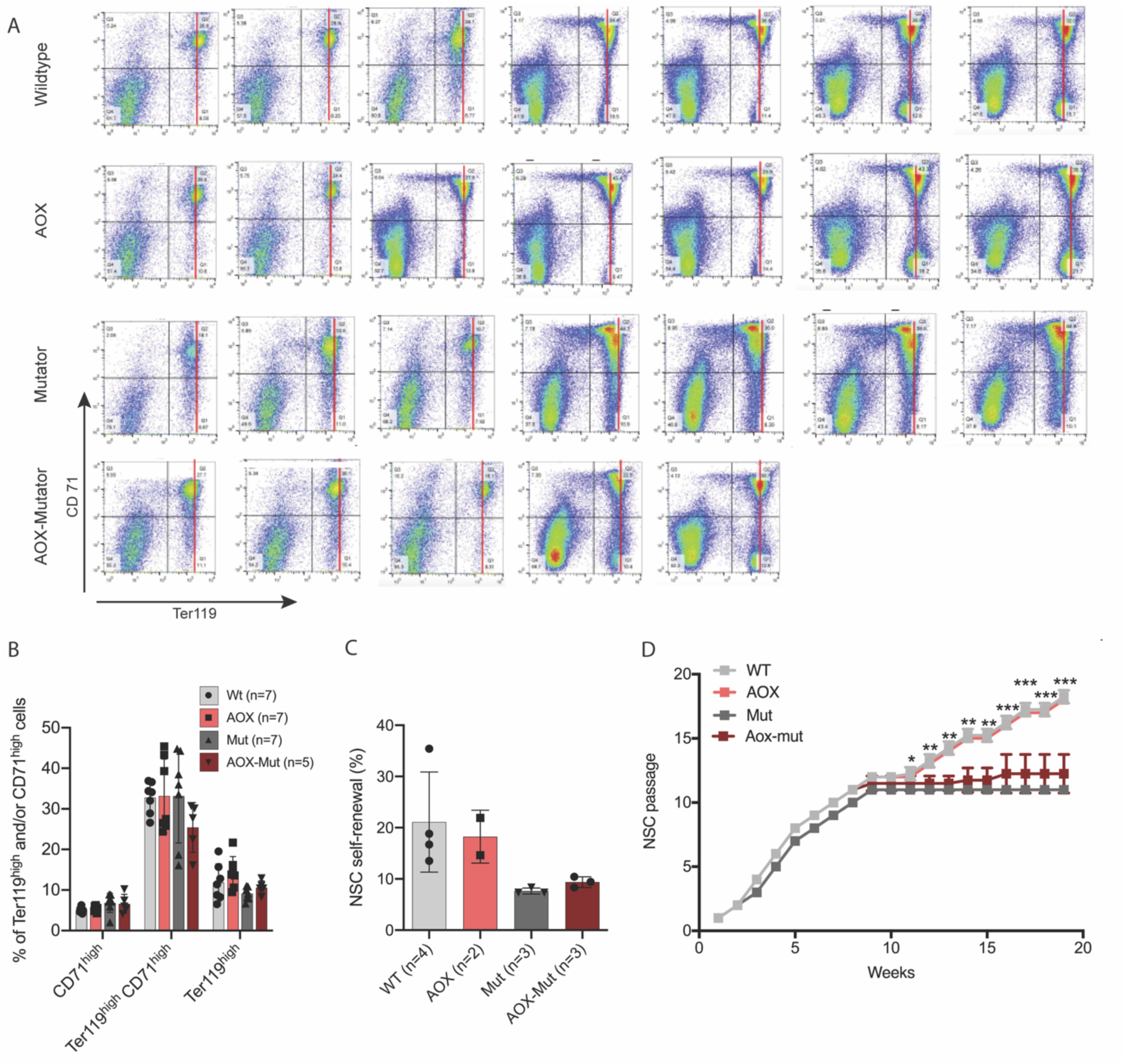
AOX shifts mutator erythropoiesis toward a wildtype state. Related to Figure 1. A) Erythrocyte precursor amounts; complete fluorescence-activated cell sorting (FACS) dot plots for Ter119 (matured) and CD71 (early-stage) populations of erythroid precursors. Black lines indicate quarters used for quantification. B) Erythrocyte precursor populations; quantification of FACS data. C) Stemness analysis of neural stem cells (NSCs) from WT, AOX, mutator and AOX-mutator mice; clonal analysis; n is number of individual mice. Altogether 10 000 – 27 000 cells per genotype were analyzed. Self-renewing cells are shown as a percentage of total cells (mean ±SD). D) NSCs growth analysis (groups same as in C). Growth defect of mutator cells compared to wildtype is significant in weeks 3–8 (****) and from week 11 onwards (shown). This is not altered by AOX expression. All graphs are mean with standard deviation (SD) with values for individual mice shown. Statistical significance determined using one-way ANOVA with p-values: * (p ≤ 0.05), ** (p ≤ 0.01), *** (p ≤ 0.001) and **** (p ≤ 0.0001).

**Figure S3.**
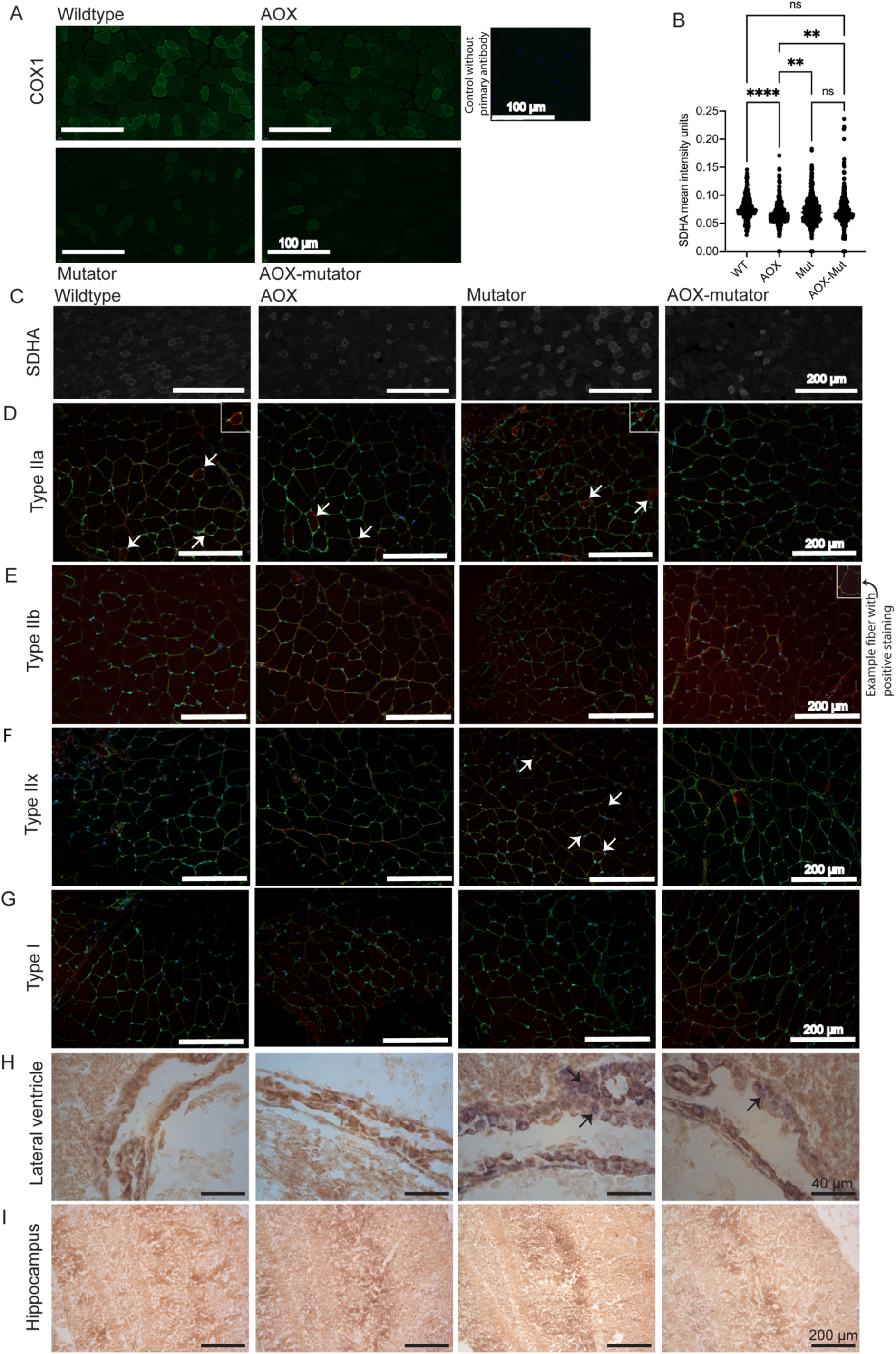
Additional histology of brain and muscle. Related to Figure 2. A) COX1 (green), protein amounts, immunofluorescence (IF) analysis. Magnification 40x, scale bar 100 µm. B) Scatterplot of SDHA mean intensity units in immunofluorescent SDHA staining in mouse skeletal muscle. C) SDHA (white), protein amounts, IF analysis. Magnification 20x, scale bar 200 µm. D–G) Myosin heavy chain (MHC) expression; immunofluorescent staining. (D) type IIa, (E) type IIb, (F) type IIx, and (G) type I in mouse skeletal muscle. Arrows indicate examples of fibers positive for MCH, stained in red. Images also include cell borders (laminin, in green) and nuclei (DAPI, in blue). Magnification 20x, scale bar 200 µm. H–I) Cytochrome *c* oxidase (COX, brown precipitate) and succinate dehydrogenase (SDH, blue precipitate) histochemical activity analysis on frozen sections; (H) lateral ventricle, subventricular zone; (I) hippocampus; of wildtype, AOX, mutator and AOX-mutator mice. Arrows show examples of fibers with SDH positivity. No abnormality in respiratory chain activities in hippocampus or dentate nucleus of mutators, but present around the lateral ventricles. H) Magnification 40x, scale bar 40 µm. I) Magnification 10 x, scale bar 200 µm. Images presented are representative samples of biological replicates. All graphs are mean with standard deviation (SD). Statistical significance determined using one-way ANOVA with p-values: * (p ≤ 0.05), ** (p ≤ 0.01), *** (p ≤ 0.001) and **** (p ≤ 0.0001).

**Figure S4.**
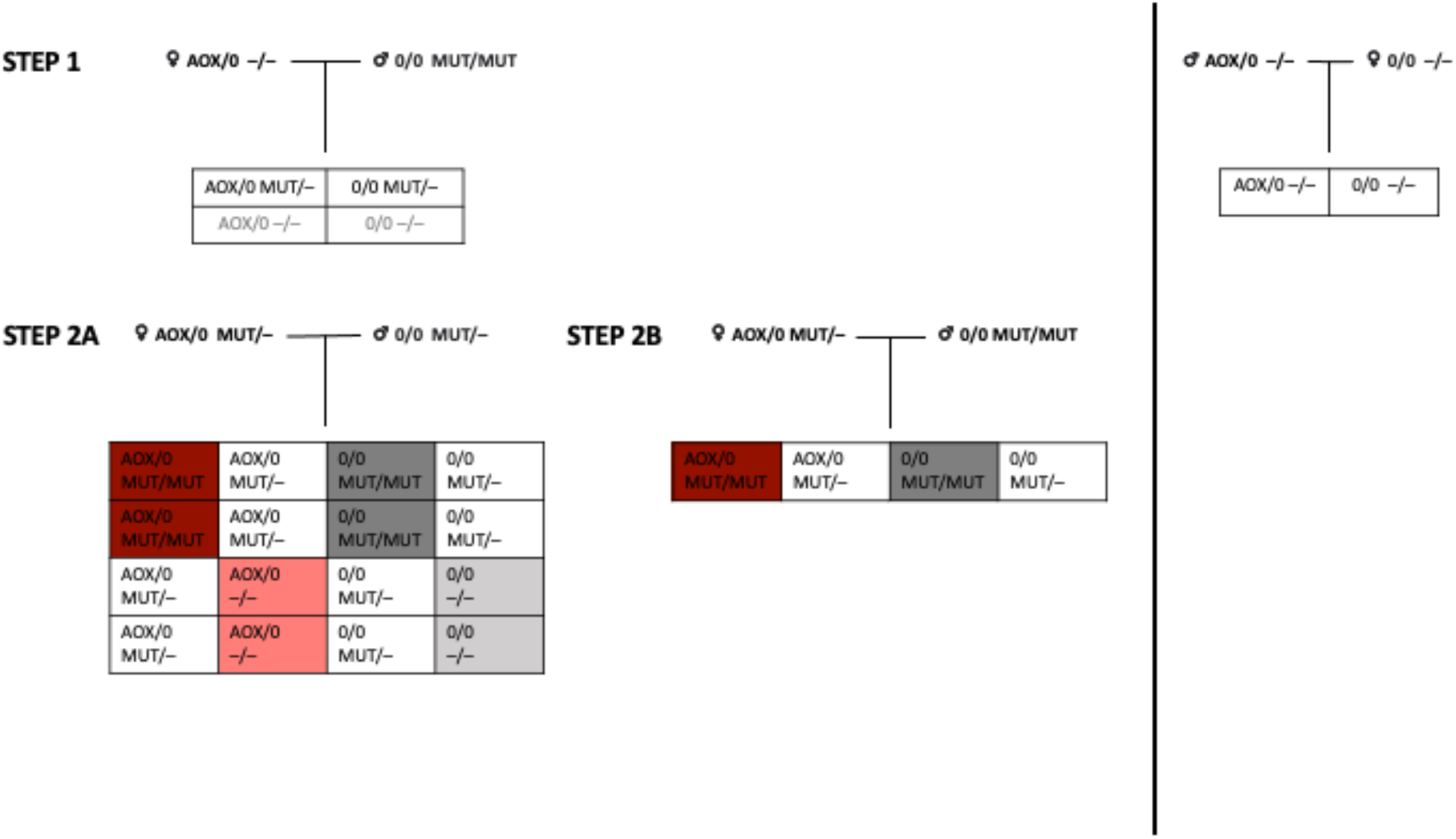
Breeding plan of AOX-mutator mice. 0 is used to mark the absence of the AOX allele and – to mark the absence of the mutator allele. Abbreviations: WT, wildtype mice; AOX, AOX mice; Mut, mutator mice; AOX-Mut, AOX-mutator mice.

